# Hierarchical Encoding of Millisecond-Scale Temporal Configuration of Sound in the Mammalian Auditory Pathway

**DOI:** 10.1101/2024.12.02.626513

**Authors:** Yuying Zhai, Peirun Song, Ana Belén Lao-Rodríguez, Xinyu Du, Haoxuan Xu, Hangting Ye, Xuehui Bao, Ishrat Mehmood, Nayaab Shahir Pandit, Cheng Chang, Zhiyi Tu, Pei Chen, Tingting Zhang, Lingling Zhang, Xuan Zhao, David Pérez-González, Manuel S. Malmierca, Xiongjie Yu

## Abstract

Natural sounds are defined not only by their spectral content but also by their fine temporal structure. Here we show that millisecond-scale temporal configuration—defined by the ordering of inter-click intervals (ICIs) within a click train—behaves as a distinct auditory feature, conserved across species and emerging hierarchically along the auditory pathway. Human listeners reliably discriminated click trains that shared the same average ICI but differed in temporal configuration, and these differences elicited robust mismatch negativity (MMN) responses in an oddball paradigm, indicating automatic cortical deviance detection. Awake rats showed analogous MMN-like ECoG responses to configuration changes, demonstrating cross-species generality. Neuropixels recordings along the inferior colliculus–medial geniculate body–primary auditory cortex (IC–MGB–A1) axis revealed minimal configuration sensitivity in IC, intermediate sensitivity in MGB, and strong stimulus-specific adaptation to temporal configuration in A1. Reversible cooling of auditory cortex reduced configuration sensitivity in MGB, implicating corticothalamic feedback in shaping thalamic representations. Layer-resolved analyses further showed that supragranular A1 neurons carry stronger configuration-specific adaptation than infragranular neurons. These findings identify temporal configuration as a feature-like dimension of sound and delineate a hierarchical, feedback-dependent IC–MGB–A1 circuit architecture for encoding fine temporal structure in the mammalian brain.

## Introduction

Sound is typically described by physical properties such as frequency, intensity, and spectral composition, which are mapped onto perceptual attributes like pitch, loudness, and timbre [1–3]. However, successful hearing in natural environments also requires the auditory system to organize acoustic input into coherent perceptual objects and streams that unfold over time [1,4,5]. This temporal organization supports the recognition of speech, music, and environmental events and underlies the distinction between “what” a sound is and “when” it occurs in complex listening scenes [2,6]. Temporal structure—the precise pattern of timing, duration, and ordering of acoustic events—is therefore a central dimension of auditory processing. Cortical activity tracks hierarchical temporal structure in connected speech, music, and narrative stimuli, reflecting integration windows that range from tens of milliseconds to many seconds [7–9]. These hierarchical temporal integration windows allow the brain to group rapidly fluctuating signals into phonemes, syllables, words, and phrases, and to distinguish regular sequences from irregular ones [10,11]. Yet despite extensive work on temporal dynamics, it remains unclear whether fine-grained, millisecond-scale temporal structure is encoded as a distinct feature of sound, separable from other acoustic dimensions.

Temporal integration provides one key mechanism by which the auditory system transforms discrete acoustic events into stable perceptual objects. When inter-click intervals (ICIs) in a click train are relatively long, listeners perceive individual clicks as separate events. As ICIs shorten below 30 ms, the clicks perceptually fuse into a continuous, pitch-like sound determined by the mean repetition rate—a phenomenon often described as temporal integration or temporal merging [12–18]. This fusion suggests that within a short ICI, the auditory system forms a unitary “temporal object” whose identity depends on the configuration of intervals, rather than on any single event. However, it is still unknown whether different millisecond-scale temporal configurations—despite having identical average ICI and spectrum—are perceived as distinct auditory features, whether such features can drive automatic deviance detection, and where along the auditory pathway a robust representation of temporal configuration first emerges.

Click trains, composed of identical brief pulses arranged over time, are therefore an attractive tool for probing temporal processing in the auditory system [17–21]. Most previous work has focused on the encoding of individual clicks and overall rate—examining temporal and rate coding as ICI varies [13,22]—rather than treating the click train as a unified temporal structure with its own identity. Building on the fusion phenomenon described above, a regular click train with constant ICI can be viewed as a single auditory object defined by its temporal configuration [15,16]. This motivates the broader hypothesis that temporal configuration—the millisecond-scale ordering of intervals within a train—acts as a distinct feature of sound, not only for regular but also for irregular click trains, and thereby contributes to sound identity and auditory object formation.

In the present study, we test this hypothesis using click trains that share the same average ICI but differ in their temporal configurations. First, we show that human listeners perceive click trains with different temporal configurations as distinct sounds, and that changes in temporal configuration elicit robust mismatch negativity (MMN) responses when served as deviants in an oddball paradigm, indicating automatic deviance detection at the cortical level. We then demonstrate that rats exhibit analogous MMN-like responses to changes in temporal configuration, suggesting a degree of evolutionary conservation in the representation of this feature. Finally, combining these behavioral and electrophysiological findings with pathway-level recordings along the rat inferior colliculus–medial geniculate body–auditory cortex (IC–MGB–A1) axis, we identify where temporal configuration sensitivity emerges along the ascending auditory pathway and provide initial insights into its underlying circuitry. Together, these results support the view that millisecond-scale temporal configuration behaves as a distinct auditory feature and provide new insight into its neural implementation across species.

## Results

### Sensitivity to Temporal Configuration in Humans

The human auditory system exhibits robust temporal integration: when the inter-click interval (ICI) in a click train is sufficiently brief, individual clicks are no longer perceived as separate events but fuse into a single auditory object. Previous work has shown that listeners typically fail to detect gaps when the ICI falls below 30 ms [17,20], a phenomenon we refer to as temporal merging. The resulting perceptual unit can be regarded as a temporally merged sound object. Here, we asked whether the temporal configuration within such temporally merged objects influences auditory perception.

We designed a protocol using ten distinct click trains (Fig 1A and S1 Table), each composed of 51 pulses with an average ICI of 4 ms. One set, labeled “Con”, maintained a constant ICI of 4 ms. Another, labeled “Asc”, featured an ascending ICI that ranged from 2 ms to 6 ms. A third set, labeled “Dec”, had a descending ICI from 6 ms to 2 ms. The set labeled “TW_10”_ included ICIs randomly selected from either 3.6 ms or 4.4 ms, representing a 10% deviation from the mean ICI of 4 ms. Additional sets labeled “TW_30”_, “TW_50”_, and “TW_70”_ included ICIs that deviated by 30%, 50%, and 70% above or below4 ms, respectively. We also created three sets (”RN_10”_, “RN_40”_, “RN_60”_) with ICIs randomly chosen within ranges of 10%, 40%, and 60% deviation from 4 ms. Because all ICIs were very short (no more than 7 ms), each train was perceived as a single temporally merged sound object; only the millisecond-scale temporal configuration differed.

**Fig 1.**
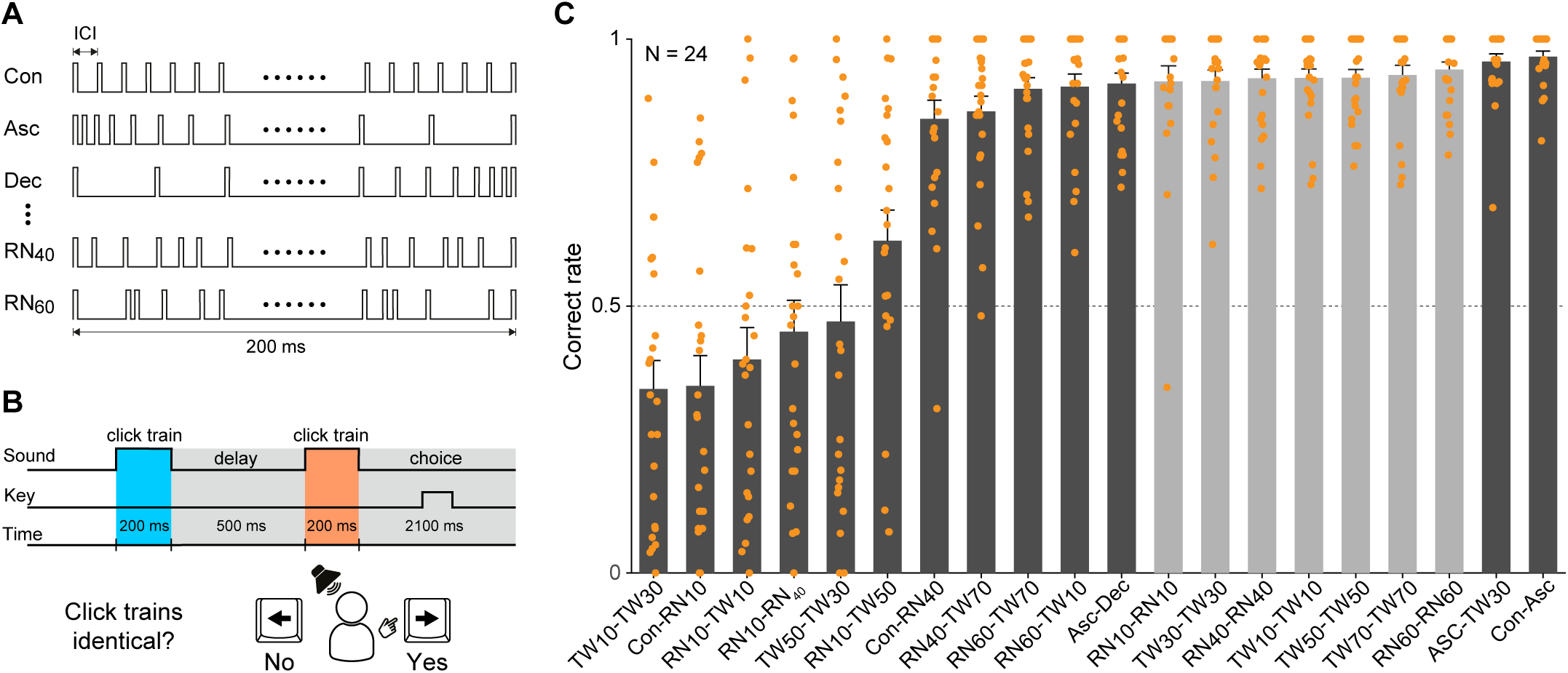
Behavioral discrimination of temporally merged click-train objects. **(A)** Different click-train temporal configurations used for the delayed match-to-sample (DMS) task using temporally merged objects. Five example sets of click trains are depicted, each comprising 51 pulses with a mean ICI of 4 ms, totaling a duration of 200 ms. Each set is labeled (e.g., Con, Asc), with extensive descriptions available in S1 Table. **(B)** Delayed match-to-sample task setup. Participants were presented with two sequences of click trains separated by a 500-ms interval. They were tasked with determining whether the two sequences were identical by pressing one of two designated buttons (left or right arrow keys). **(C)** Behavioral performance across participants. Response accuracy is displayed as the mean correct response rate for each tested click train pair. Dark grey bars represent trials with “different” pairs, and light gray bars represent trials with “identical” pairs. Error bars indicate the standard error of the mean (SEM). Orange circles denote individual participant accuracy (N = 24 participants).

To assess perceptual sensitivity to temporal configuration, we used a delayed match-to-sample task in which subjects heard two 200-ms click trains separated by a 500-ms silent interval (Fig 1B). On each trial, the two trains were either identical or drawn from a specific configuration pair, and participants reported whether the two sounds were the “same” or “different” via button press. Twenty-four participants engaged in the task involving 20 pairs of click trains (Fig 1C), and they exhibited similar psychological response patterns characterized by poor discrimination between the TW_10-_TW_30,_ Con-RN_10,_ and RN_10-_TW_10 p_airs and excellent discrimination between Con-Asc, Asc-TW_30,_ Asc-Dec, RN_60-_TW_10 a_nd RN_60-_TW_70 p_airs. On average, the correct response rates for the TW_10-_TW_30,_ Con-RN_10 a_nd RN_10-_TW_10 p_airs were 0.34, 0.35, and 0.40 (Fig 1C), respectively, all significantly below the chance level of 0.5 (p < 0.05 for the three pairs, Kolmogorov-Smirnov test). In contrast, the correct response rates for the Con-Asc, Asc-TW_30,_ Asc-Dec, RN_60-_TW_10 a_nd RN_60-_TW_70 p_airs were 0.967, 0.958, 0.921, 0.921 and 0.907 (Fig 1C), respectively, indicating that participants recognized the two temporally merged auditory objects as distinct entities. When both click trains were identical, the correct response rates were also very high (between 0.911 and 0.933). These behavioral results demonstrate that, even when click trains are matched in average ICI and duration, differences in temporal configuration can be perceived as differences between distinct auditory objects. The graded pattern of discrimination across configuration pairs suggests that temporal configuration is a candidate feature for sound identification, motivating a search for neural correlates.

We next constructed an oddball paradigm using TW30 as the standard sound and TW50, Asc, RN40 and TW10 etc. as the deviant sounds (Fig 2A). The deviant sounds were interspersed randomly among repetitive standard sounds, with a pattern of average nine standard sounds followed by one deviant sound at 300-ms ICI. We conducted 64-channel electroencephalogram (EEG) recordings (Fig 2B) on 32 subjects to capture the neural responses to these auditory stimuli. In one illustrative case, the deviant response to TW_50 c_ontrasted sharply with the response to the preceding standard sound TW_30._ Specifically, robust responses were detected to TW_50,_ particularly in the occipital and temporal lobes, whereas no obvious response was observed for the preceding standard sound TW_30 (_Fig 2B). An enlarged view of channel TP8 clearly demonstrated the difference (Fig 2C, left panel), with alignments based on the onsets of the sounds (Fig 2C, right panel). Grouping 27 channels in the occipital and temporal lobes (indicated by red background in Fig 2B) further clarified the difference between deviant and standard responses, revealing a sustained and statistically significant contrast almost from 80 ms to 280 ms (p < 0.05, deviant vs. standard, independent-samples t-test, with FDR correction applied, Fig 2D).

**Fig 2.**
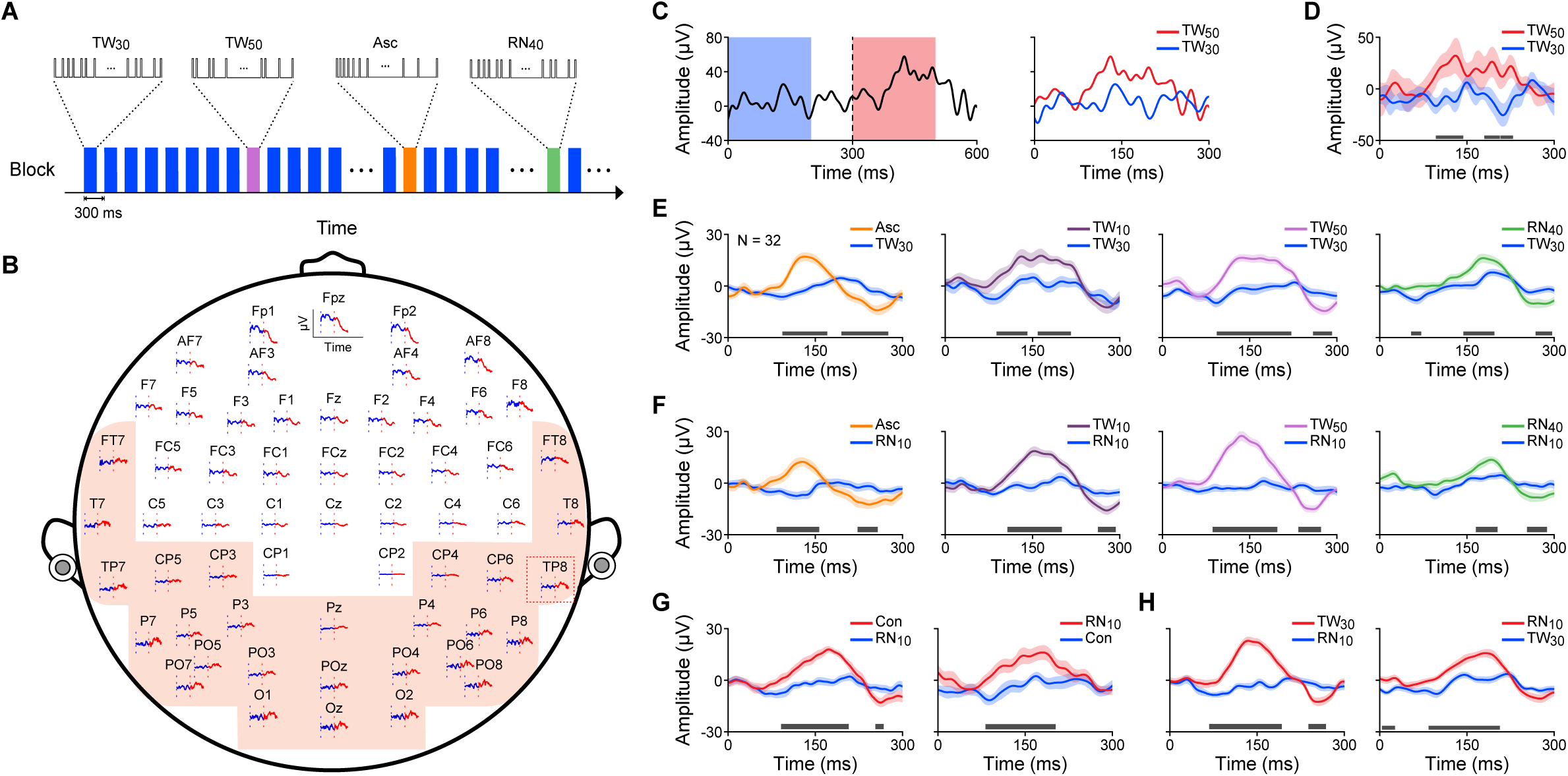
EEG response to oddball paradigm with temporally merged objects in humans. **(A)** Oddball Paradigm Setup: the standard sound (TW_30)_ is presented at a 300-ms inter-stimulus interval, followed by a deviant sound after 7-11 standard repetitions. The deviant sound is randomly chosen from a set of stimuli (TW_50,_ Asc, RN_40,_ etc.). **(B)** Topographical layout of the 64-channel EEG setup used for recording human auditory responses. For each channel, individual EEG traces are shown in response to the standard sound (TW_30,_ blue) and the subsequent deviant sound (TW_50,_ red). Two vertical dotted lines mark the onset of the standard sound (preceding the deviant, blue line) and the onset of the deviant sound (red line) respectively. Shaded background indicates channels that showed robust auditory responses and were selected for further analysis. The red rectangle indicates the channel expanded in panel (C). **(C)** Left panel: Enlarged view of the EEG response from the channel indicated by the red rectangle in (B). Right panel: Comparative analysis of the EEG responses to the standard and deviant sounds, with alignments based on the onsets of the sounds. **(D)** The averaged responses to the standard and deviant sounds across channels highlighted with the shaded background, for the example participant. A grey bar above the X-axis indicates a significant difference (p<0.05, independent-samples t-test with FDR correction) between the standard and deviant responses during this period. **(E)** Population-level EEG responses (N = 32 subjects) to the standard sound (TW_30,_ blue) and four different deviants (Asc, TW_10,_ TW_50 a_nd RN_40)_. A grey bar above the X-axis indicates a significant difference (p<0.05, independent-samples t-test with FDR correction) between the standard and deviant responses during this period. **(F)** Population-level EEG responses to the standard (RN_10,_ blue) and four different deviants (Asc, TW_10,_ TW_50 a_nd RN_40)_. **(G)** Population-level EEG responses to the stimulus pair Con-RN_10,_ when each of them is the standard (blue) or deviant (red) in a classic oddball paradigm. **(H)** Population-level EEG responses to the stimulus pair TW_30-_RN_10,_ when each of them is the standard (blue) or deviant (red) in a classic oddball paradigm.

In our human subject population, significant differences emerged when the deviants were Asc, TW_10,_ TW_50,_ RN_40,_ etc. particularly within the window mainly from 100 ms to 200 ms. Fig 2E and 2F show examples of such differences when the standard was TW_30 (_Fig 2E) and RN_10 (_Fig 2F), where the grey horizontal bars indicate time ranges where the response to the deviant was significantly different from the response to the standard (p < 0.05, independent-samples t-test, with FDR correction applied). Additional stimulus pairs also showed significant differences. For example, significant MMN responses were observed when RN_10 a_nd Con were paired as the standard and deviant (Fig 2G), and likewise when RN_10 a_nd TW_30 w_ere paired as the standard and deviant (Fig 2H). Thus, the human cortex automatically detects changes in temporal configuration, generating MMN responses even in the absence of an explicit discrimination task. These findings support the view that temporal configuration is encoded as a basic sound feature at the cortical level.

### Cross-Species Sensitivity to Temporal Configuration in Rats

Building on the psychological and electrophysiological evidence supporting the role of temporal configuration as a distinct sound feature in humans, we expanded our research to assess whether this phenomenon is conserved across species. Specifically, we conducted experiments on awake rats to determine if similar auditory processing mechanisms are evident. We performed 32-channel electrocorticography (ECoG) recordings of the auditory cortex in awake rats (Fig 3A), using stimuli analogous to those used in the human experiments (Fig 2A).

**Fig 3.**
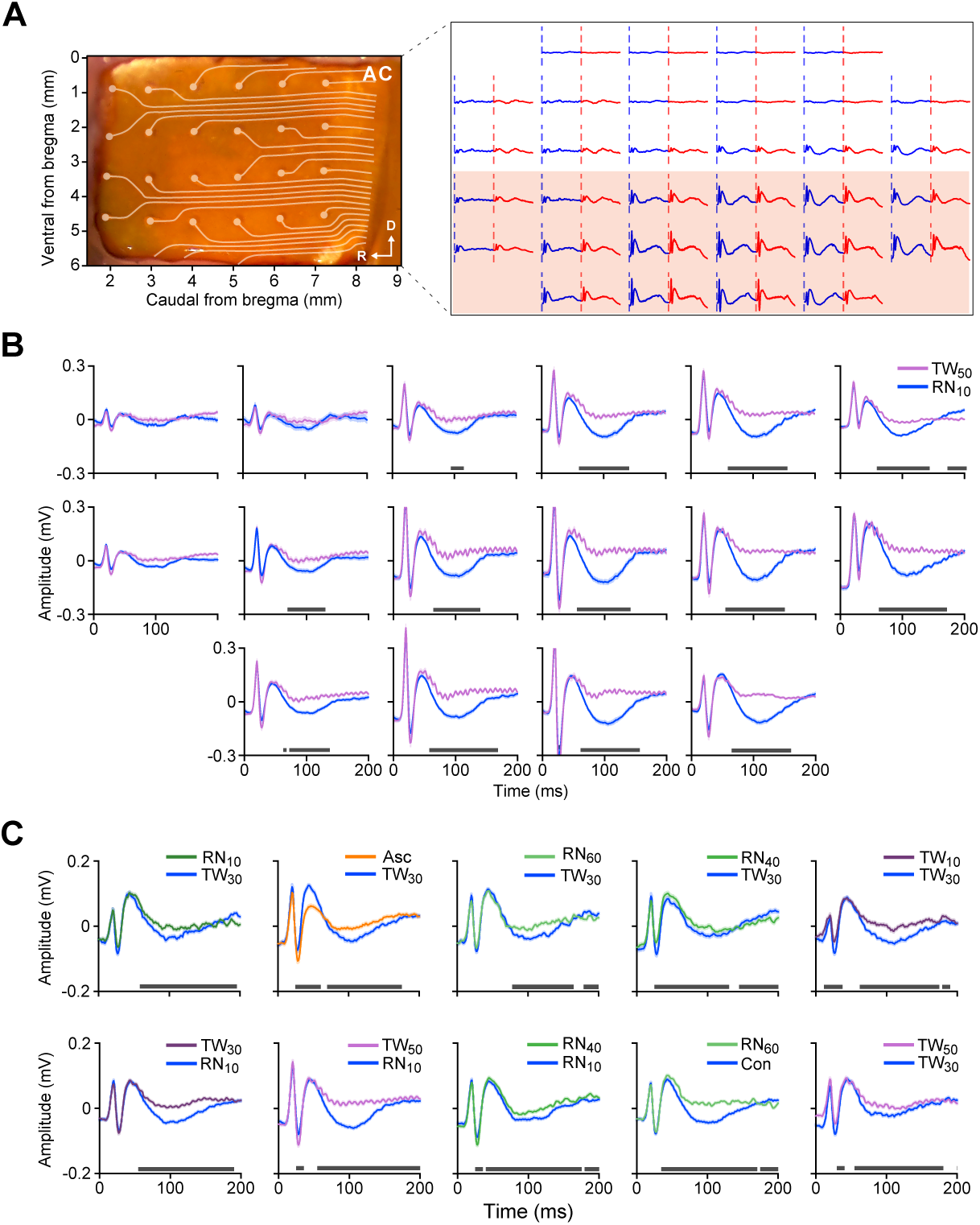
ECoG response to oddball paradigm with temporally merged objects in rats. **(A)** ECoG Recording Setup: Left panel shows a schematic a picture of the craniotomy over the AC, overlaid with the 32-channel ECoG setup utilized for recording auditory responses in rats. Right panel displays the responses of the 32-channel ECoG to one standard sound and the subsequent deviant sound. Two vertical dotted lines mark the onset of the standard sound (preceding the deviant, blue line) and the onset of the deviant sound (red line) respectively. Shaded background indicates channels that exhibited robust auditory responses, selected for further detailed analysis. **(B)** Responses to the standard (RN_10)_ and subsequent deviant sounds (TW_50)_ in the channels outlined by the red rectangle, with alignments based on the onsets of the sounds, for one example rat. **(C)** Population-level ECoG responses to the standard sound and its subsequent deviant sounds across ten stimulus pairs. The top row shows responses to TW_30 a_s standard (blue) and RN_10,_ Asc, RN_60,_ RN_40 a_nd TW_10 a_s deviants (different colors). The bottom row shows responses to RN_10 a_s standard (blue) and TW_30,_ TW_50 a_nd RN_40 a_s deviants (different colors), Con-RN_60 (_standard and deviant, blue and green, respectively) and TW_30-_TW_50 (_standard and deviant, blue and purple, respectively). ECoG channels in each rat were selected based on deviant responses exceeding the baseline amplitude ([-50, 0 ms]) by more than 2 standard deviations within the [0, 200 ms] window (N = 3 rats, n = 36 channels). Similarly, TW_30-_RN_10 a_nd RN_10-_TW_30 (_left column) constitute a classic oddball paradigm. A bar above the X-axis indicates a significant difference (p<0.05, independent-samples t-test with FDR correction) between the standard and deviant responses during this period.

Significant differences between standard and deviant click trains were detected in most channels eliciting auditory responses. For instance, Fig 3B illustrates the auditory responses of an example rat to RN_10 a_nd TW_50 c_lick trains across multiple channels. To summarize across rats, we averaged the significant channels (see Methods) across all rats. The results revealed widespread significant differences between standard and deviant sounds in other scenarios as well, where both the standard and deviant sounds varied (Fig 3C). These data indicate that, like humans, rats exhibit MMN-like responses to changes in temporal configuration of click trains, despite identical average ICIs and overall energy. The cross-species consistency suggests that temporal configuration is a fundamental dimension of auditory processing that is preserved across evolution and can be studied mechanistically in animal models.

### Neuronal Mechanisms Underlying Sensitivity to Temporal Configuration

Next, we conducted extracellular recordings along the auditory pathway to further investigate the underlying neuronal mechanisms (Fig 4A). These recordings include A1 (Fig 4C), medial geniculate body (MGB, Fig 4D), and inferior colliculus (IC, Fig 4E). This approach aimed to track the origin of sensitivity to click-trains temporal configurations using an oddball paradigm, a widely adopted method for assessing stimulus-specific adaptation at the neuronal level [23–34] (Fig 4B). The oddball paradigm comprised two sets, each with a pair of temporally merged click trains. In one set, we repeated one train (standard train) nine times followed by the other train (deviant) and then repeated this pattern composed of nine standards and one deviant for 30 cycles; in the other set, we reversed the standard and deviant trains. For comparative analysis, the response to the standard sound immediately preceding the deviant sound was selected as standard response.

**Fig 4.**
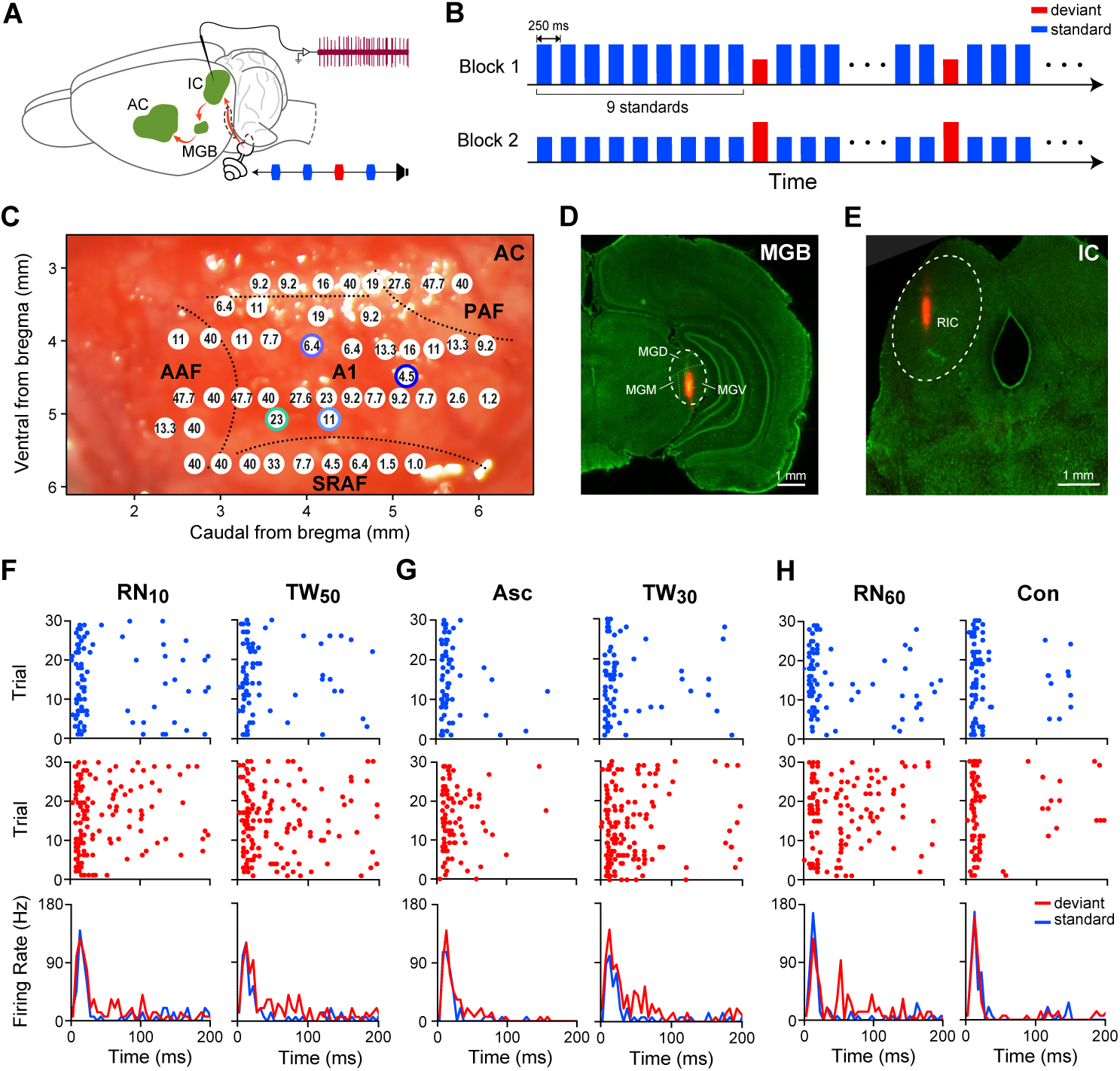
Neuronal response to oddball paradigm with temporally merged objects. **(A)** Extracellular recording setup: the recording process along the auditory pathway, including the inferior colliculus (IC), medial geniculate body (MGB), and primary auditory cortex (A1). Modified from Parras et al. (2017). **(B)** Oddball Stimulus Paradigm: the oddball paradigm consists of two sets of stimuli. Each set contains a pair of sounds: one sound is repeatedly presented as the standard sound at a 250-ms inter-stimulus interval, followed by a deviant sound after every nine standard sounds. This sequence (9 standard sounds followed by 1 deviant) is repeated 30 times. In the other set, the roles of the standard and deviant sounds are reversed. **(C)** Tonotopic distribution of characteristic frequencies (CF) across the auditory cortex of one example rat, with frequencies indicated in kilohertz (kHz). Colored circles mark the locations where neuronal recordings were conducted for the main dataset in A1 in the rat. **(D)** Localization of the MGB. The orange area highlights the tracking of Neuropixel probe within the MGB, as indicated by the thick dashed line. The thin dashed lines delineate the subregions of the MGB, including the medial (MGM), dorsal (MGD), and ventral (MGV) subdivisions. **(E)** Localization of the IC. The orange area indicates the tracking of the Neuropixel probe within the IC, outlined by the thick dashed line. This particular electrode was placed in the rostral subdivision of the inferior colliculus (RIC). **(F-H)** Responses of an example A1 neuron in the oddball paradigm. Raster plots display the responses to a sound when it is in the role of standard (top row) and in the role of deviant (middle row), for three stimulus pairs: **(F)** RN_10 a_nd TW_50;_ **(G)** Asc and TW_30;_ **(H)** RN_60 a_nd Con. The bottom row presents the peri-stimulus time histograms (PSTHs) for each sound when acted as deviant (red) or standard (blue) stimulus.

For one example A1 neuron, we tested three pairs: RN_10 a_nd TW_50,_ Asc and TW_30,_ and RN_60 a_nd Con. In the pair RN_10 a_nd TW_50,_ the A1 neuron demonstrated robust responses when the trains were presented as deviants (raster plots in red, Fig 4F), with significant differences between deviant and standard responses observed for both trains (RN_10,_ p = 0.0014; TW_50,_ p<0.001; independent-samples t-test, bottom row in Fig 4F). Similarly, for the pair Asc and TW_30,_ we observed analogous responses, with significant differences between deviant and standard responses for both (Asc, p < 0.05; TW_30,_ p<0.001; independent-samples t-test, bottom row in Fig 4G). For the pair RN_60 a_nd Con, significant differences were observed between deviant and standard responses for RN_60 (_p = 0.0102, independent-samples t-test, bottom row in Fig 4H) but not for Con (p = 0.94, independent-samples t-test, bottom row in Fig 4H).

We further analyzed the differences in neuronal responses across different levels of the auditory pathway for the pairs RN_10 a_nd TW_50,_ RN_60 a_nd Con, and Asc and TW_30 (_Fig 5), presented in an oddball paradigm. For RN_10 a_nd TW_50,_ the responses in the IC neuronal population (n = 940) to deviant and standard sounds overlapped and showed no significant differences (RN_10,_ p = 0.81; TW_50,_ p = 0.18; independent-samples t-test, Fig 5C). However, in the MGB (n = 632), deviant responses for both RN_10 a_nd TW_50 w_ere significantly stronger than standard responses (RN_10,_ p = 0.018; TW_50,_ p = 0.0019; independent-samples t-test, Fig 5B). Similarly, in the A1 (n = 671), deviant responses were significantly stronger than standard responses (RN_10,_ p = 0.037; TW_50,_ p < 0.001; independent-samples t-test, Fig 5A).

**Fig 5.**
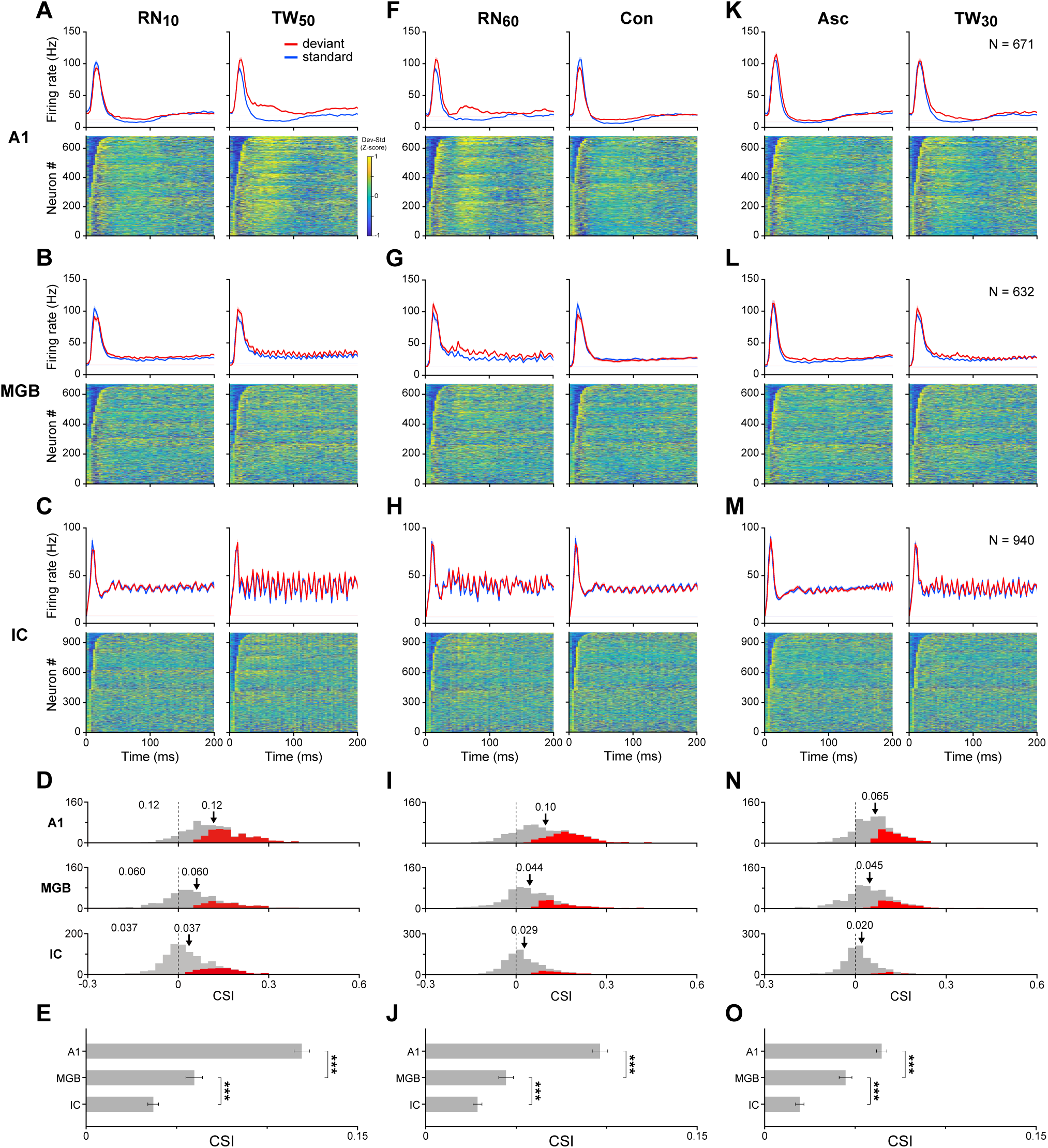
Populational neuronal response to oddball paradigm with temporally merged objects in IC, MGB, and A1. **(A)** Top panels: Population PSTHs displaying averaged neuronal responses to RN_10 (_left) and TW_50 (_right) as deviant (red) and standard (blue) stimuli in A1. Bottom panels: Heatmaps illustrating the z-scored differences (deviant - standard) in PSTHs of firing rates for each neuron across the population. Warmer colors indicate higher positive differences, while cooler colors indicate negative differences. Time is aligned to stimulus onset at 0 ms. **(B-C)** Population PSTHs in MGB and IC following the same formatting conventions as A1 (A). **(D)** Distribution of CSI calculated for neurons in A1 (top row), MGB (middle row), and IC (bottom row) within the time window of 5-200 ms from stimulus onset. Red bars highlight neurons with CSI significant greater than 0, determined by a nonparametric bootstrap procedure with 1,000 iterations (p < 0.05). Arrows point to the mean. **(E)** Comparison of CSI values among A1, MGB, and IC. Error bars represent standard error of the mean (SEM). ***: p<0.001, ANOVA with post-hoc test. **(F-J)** Population responses to another sound pair in the oddball paradigm (RN_60,_ Con), following the same conventions as panels (A-E). **(K-O)** Population responses to yet another sound pair in the oddball paradigm (Asc, TW_30)_, adhering to the same conventions as panels (A-E).

To further characterize the differences between standard and deviant responses across the auditory pathway, we employed the common stimulus-specific adaptation index (CSI). This index is defined as the sum of the differences between the deviant and standard responses for the two trains, normalized by the sum of both the deviant and standard responses for both trains (see Methods). The analysis showed that the distribution of the CSI shifted positively (Fig 5D), indicating a greater distinction between deviant and standard responses as auditory processing progresses from the IC to A1. The average CSI increased significantly from the IC to A1 (p < 0.001 for both comparisons: A1 vs. MGB and MGB vs. IC, ANOVA with post-hoc test, Fig 5E).

Consistent sensitivity to deviant sounds was observed across different sound pairs in the population. For RN_60 a_nd Con, differences between deviant and standard responses were minimal in the IC (RN_60,_ p = 0.31; Con, p = 0.90; independent**-**samples t-test, Fig 5H) but became significantly pronounced in A1 (RN_60,_ p < 0.001; Con, p < 0.001; independent**-**samples t-test, Fig 5F). The CSI distribution similarly shifted positively (Fig 5I), with the average CSI showing a significant increase from IC to A1 (p < 0.001 for both comparisons: A1 vs. MGB and MGB vs. IC, ANOVA with post-hoc test, Fig 5J). For Asc and TW_30,_ differences between deviant and standard responses were minimal in the IC (Asc, p = 0.71; TW_30,_ p = 0.42; independent**-**samples t-test, Fig 5M) but became significantly pronounced in A1 (Asc, p = 0.035; TW_30,_ p < 0.001; independent**-**samples t-test, Fig 5K). The CSI distribution similarly shifted positively (Fig 5N), with the average CSI showing a significant increase from IC to A1 (p < 0.001 for both comparisons: A1 vs. MGB and MGB vs. IC, ANOVA with post-hoc test, Fig 5O). The results from the subdivisions of MGB and IC are summarized in S1 Fig.

The relatively lower CSI with click trains in the IC may stem from the fact that IC neurons usually exhibit less SSA compared to neurons in the MGB and A1 [24,28,33,35]. To rule out this possibility, we specifically selected neurons that demonstrated stronger deviant responses compared to the standard responses (i.e., CSI significantly greater than 0) for a pair of two complex tones (CT, see Method, *Session 5 Complex Tone Oddball procedure*) in an oddball paradigm, from A1 (n = 94), MGB (n = 109), and IC (n = 140) (S2 Fig). In this dataset, CSI was notably higher in the IC than in A1 (p < 0.001, Wilcoxon rank-sum test, S2B-C Fig). However, using the same dataset with the three pairs of click trains, the CSI still tended to increase from IC to A1 (S2E, H and K Fig), and remained significantly higher in A1 than in IC for the three pairs (p<0.001, ANOVA with post-hoc test t-test, S2F, I and L Fig). Thus, the low CSI for click trains in IC reflects a reduced ability to discriminate temporal configurations, not a general lack of SSA capacity, whereas the higher CSI in A1 reflects enhanced configuration discrimination rather than a generic increase in SSA.

Since MGB and AC are extensively connected [36], the sensitivity to temporal configuration in MGB may arise from AC top-down inputs. To explore this possibility, we used cooling techniques to decrease neuronal activities in the auditory cortex (S2 Fig) and thus decrease the top-down inputs to MGB. In the normal condition before cooling the AC, the MGB neuronal population (n = 86) demonstrated robust SSA for the three pairs, and the deviant responses were stronger than the standard responses (RN_10,_ p = 0.18; TW_50,_ p < 0.001, top row in Fig 6A; RN_60,_ p < 0.001; Con, p = 0.40, top row in Fig 6D; Asc, p < 0.05; TW_30,_ p < 0.001, top row in Fig 6G; independent-samples t-test) with average CSIs of 0.12, 0.13, and 0.089 respectively (top row in Fig 6B, 6E and 6H). During the cooling of AC, the same MGB population demonstrated similar responses between deviant and standard responses for both pairs (RN_10,_ p = 0.86; TW_50,_ p = 0.59, bottom row in Fig 6A; RN_60,_ p = 0.29; Con, p = 0.99, bottom row in Fig 6D; Asc, p = 0.98; TW_30,_ p = 0.50, bottom row in Fig 6G; independent-samples t-test) with average CSIs of 0.038, 0.079 and 0.071 respectively (bottom row in Fig 6B, E and H). Three sessions of experiments were carried out involving three rats. The CSI of the cooling condition was significantly smaller than the normal pre-cooling condition for the pair of RN_10 a_nd TW_50 (_p < 0.001, Fig 6C, paired t-test), and the pair of RN_60 a_nd Con (p=0.0019, Fig 6F, paired t-test). However, for the pair of Asc and TW_30,_ the CSIs were similar between the cooling condition and the normal pre-cooling condition (p=0.095, Fig 6I, paired t-test), mainly due to the fact that the CSI is 0.089 in the normal condition, which is already very small. Thus, the sensitivity to temporal configuration in MGB probably arises from the top-down input from AC.

**Fig 6.**
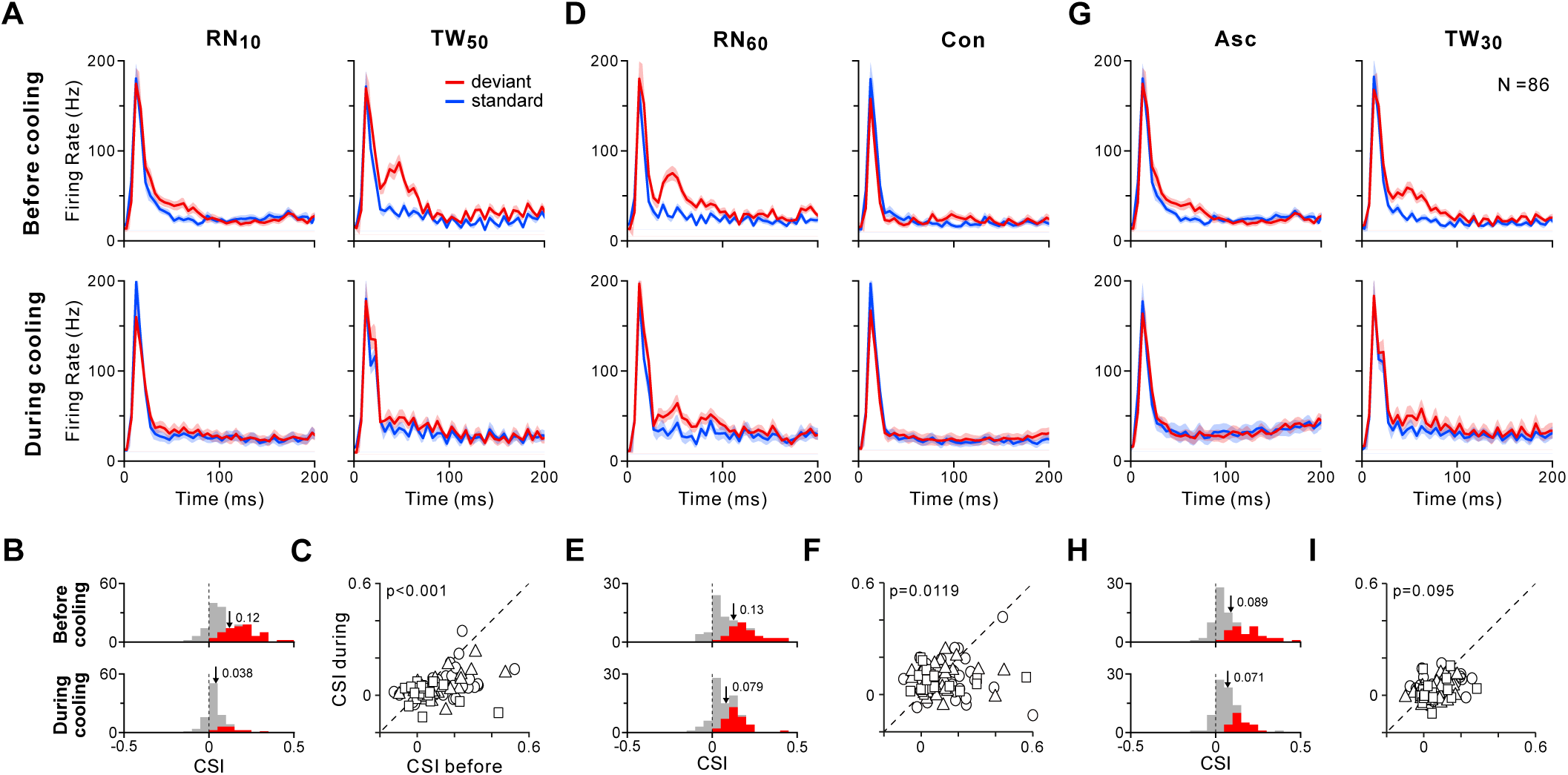
Effect of AC cooling on the response to oddball paradigm with temporally merged objects in MGB. **(A)** Populational PSTHs displaying responses to RN_10 (_left) and TW_50 (_right) as deviant (red) and standard (blue) stimuli under two conditions: before AC cooling (top row), and during AC cooling (bottom row). **(B)** Distribution of CSI before cooling (top row) and during cooling (bottom row), with arrows indicating the means. Red bars highlight neurons with CSI significant greater than 0, determined by a nonparametric bootstrap procedure with 1,000 iterations (p < 0.05). **(C)** Comparison of CSI between before and during AC cooling with different signs representing data from different rats (N = 3 rats, total neurons = 86). p < 0.001, paired t-test. **(D-F)** Populational responses from the same dataset to the sound pair in the oddball paradigm (RN_60,_ Con), following the same conventions as panels (A-C). p=0.0119, paired t-test. **(G-I)** Populational responses to sound pair in the oddball paradigm (Asc, TW_30)_, adhering to the same conventions as panels (A-C). p=0.095, paired t-test.

Since AC plays a crucial role in sensitivity to temporal configuration, we next asked which cortical layers are most involved in configuration coding. Using linear array recordings with 384-channel Neuropixel electrodes, we categorized the six cortical layers into three groups based on the current source density (CSD) profile (S4 Fig): the supra layer (Ⅰ/Ⅱ, n = 30), middle layer (Ⅲ/Ⅳ, n = 176), and deep layer (Ⅴ/Ⅵ, n = 465). For the same neuronal population, the average CSI values for the pair RN_10 a_nd TW_50 w_ere 0.162, 0.166, and 0.099 in the supra, middle, and deep layers, respectively (Fig 7A). For the pair RN_60 a_nd Con, the average CSI values were 0.145, 0.136, and 0.08 (Fig 7C), and for the pair Asc and TW_30,_ the average CSI values were 0.106, 0.090, and 0.052 (Fig 7E), respectively. When analyzing complex tones, the average CSI values were 0.073, 0.11, and 0.053 (Fig 7G) across the supra, middle, and deep layers. Interestingly, the CSI in the supra layer was significantly greater than in the deep layer (p < 0.001, ANOVA with post-hoc test) for all three pairs of click trains, but similar between layers for complex tones (p = 0.45, ANOVA with post-hoc test). These findings suggest that superficial cortical layers play a particularly important role in encoding temporal configuration, whereas layer-specific differences are less marked for frequency-related features.

**Fig 7.**
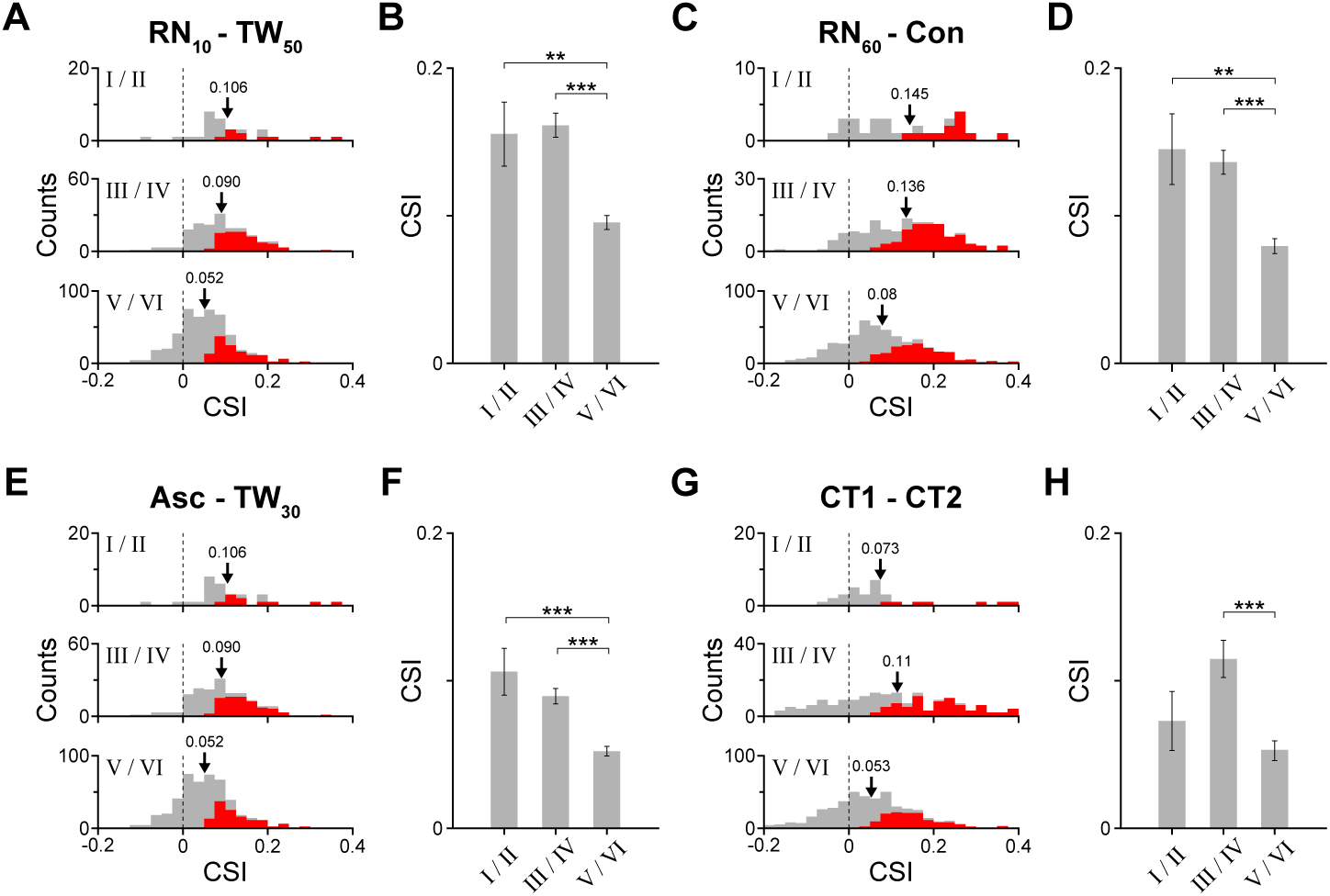
Neuronal responses to temporally merged objects and complex tones across cortical layers in A1 in the oddball paradigm. **(A)** Distribution of CSI in the oddball paradigm (RN_10,_ TW_50)_ across cortical layers: supra layer (layers Ⅰ/Ⅱ, n = 30, top row), middle layer (layers Ⅲ/Ⅳ, n = 176, middle row), and deep layer (layers Ⅴ/Ⅵ, n = 465, bottom row). Arrows point to the mean. **(B)** Comparison of CSI among supra, middle, and deep layers. Red bars highlight neurons with CSI significant greater than 0, determined by a nonparametric bootstrap procedure with 1,000 iterations (p < 0.05). Error bars represent standard error; n.s., no significant difference; **, p < 0.01; ***, p < 0.001, ANOVA with post-hoc test. **(C-H)** Populational responses from the same datasets featured in panels (A-B) to two additional sound pairs: (C-D) RN_60 a_nd Con, (E-F): Asc and TW_30,_ and complex tones pair (G-H). Following the same conventions as panels (A-C).

## Discussion

In this study, we demonstrate that millisecond-scale temporal configuration is a distinct auditory feature, conserved across species and expressed hierarchically along the auditory pathway. In humans, listeners reliably discriminated click trains that differed only in temporal configuration (Asc, Dec, TW_10/30/50/70,_ RN_10/40/60)_, revealing robust perceptual sensitivity to fine temporal structure (Fig 1). Changes in configuration elicited clear mismatch negativity (MMN) responses in EEG (Fig 2), indicating automatic cortical deviance detection. In rats, analogous MMN-like responses were observed with ECoG (Fig 3), suggesting evolutionary conservation of configuration-based processing. Neuropixels recordings along the IC–MGB–A1 axis showed weak configuration sensitivity in IC, intermediate sensitivity in MGB, and strong stimulus-specific adaptation in A1 (Fig 5 and S2 Fig), consistent with a hierarchical emergence of configuration encoding. Reversible cooling of auditory cortex reduced configuration sensitivity in MGB (Fig 6), demonstrating that thalamic representations are shaped by corticothalamic feedback. Layer-specific analyses further revealed higher SSA indices for temporal configuration in supragranular than infragranular layers of A1 (Fig 7), implicating superficial cortical circuits in coding fine temporal structure. Together, these findings identify temporal configuration as a fundamental, feature-like dimension of hearing—on a par with frequency or intensity—implemented by a hierarchical, feedback-dependent cortical–thalamic network.

### Temporal Configuration as a New Feature of Sound

Sound is commonly characterized by parameters such as frequency, intensity, duration, timbre, and spatial location. However, the temporal dimension of auditory processing is equally fundamental. The brain automatically tracks temporal regularities over hundreds of milliseconds, often generating sustained neural responses to structured sound sequences [10,11,37]. By contrast, how the auditory system encodes temporal structure at the millisecond scale remains much less understood. Temporal configurations are ubiquitous in nature, manifesting in the fine timing of water flow, bird songs, insect calls, and wind in trees.

Advances in neuroimaging and electrophysiology have revealed hierarchical temporal integration for spoken language in humans [7,9], but there is a substantial gap in our understanding of how short-timescale temporal configuration is represented in non-human animals and in the underlying neuronal circuitry. In this context, click trains offer a particularly useful approach, because they allow temporal configuration to be manipulated while largely controlling other acoustic dimensions. Historically, click trains have revealed rich neuronal responses in the auditory system, but most work has focused on how individual clicks or average rate are encoded rather than on the configuration of the sequence as a whole [13,19,38]. When the ICI falls below 30 ms, regular click trains are generally perceived as a single pitch-like sound [12], consistent with temporal merging. What has remained unclear is whether different temporal configurations within such merged objects are treated as distinct auditory features.

Our data directly address this question. In a delayed match-to-sample task, participants reliably discriminated click trains that differed only in temporal configuration, with performance showing a graded pattern across configuration pairs (Fig 1). This robust and consistent performance suggests that temporal configuration is a salient perceptual dimension. Moreover, oddball paradigms using pairs such as Asc–TW_30,_ TW_10–_TW_30,_ TW_50–_TW_30,_ and RN_40–_TW_30 e_licited clear MMN responses (Fig 2E), indicating that the brain automatically differentiates these configurations and represents them as distinct auditory objects even without an explicit task. These findings underscore the profound impact that temporal configuration can have on auditory perception and extend our understanding of how the auditory system interprets complex sounds.

Importantly, our findings are not limited to humans but extend across species. Both human and rat data showed similar MMN/MMN-like signatures for changes in temporal configuration (Figs 2 and 3), suggesting that configuration-based processing is a general property of mammalian auditory systems rather than a specialization for speech or music. This cross-species consistency not only highlights the fundamental nature of temporal processing in auditory perception but also informs our understanding of auditory ecology and the evolutionary pressures that shape sensory systems [39]. Recognizing temporal configuration as a distinct feature of sound opens new avenues for basic and applied research, including the design of auditory signal-processing strategies and devices that exploit temporal structure more effectively.

### Neuronal Sensitivity to Click-Train Temporal Configuration along the Auditory Pathway

We examined neuronal sensitivity to temporal configuration using SSA within an oddball paradigm, assessing responses across multiple levels of the auditory pathway, from IC to A1 (Fig 4A). SSA has been extensively studied in the auditory cortex [31,35,40–47], medial geniculate body [42,48–53], inferior colliculus [32,54–58], and cochlear nucleus [30,59]. Typically, these studies have utilized pure tones to generate oddball stimuli, with previous data suggesting that SSA is mediated by adaptation in narrowly-tuned inputs, a concept supported by computational modeling [60] and the discovery of synaptic SSA [26]. However, recent studies propose that SSA is predominantly a predictive process [4,33,61–63]. Our paradigm provides a stringent test of these ideas because all click trains are constructed from identical pulses and share the same average ICI; spectral content and overall rate are held constant, and only temporal configuration differs. Under such conditions, simple feedforward adaptation would be expected to produce similar adaptation across configurations. Consistent with this, we observed minimal SSA in IC and only modest SSA in MGB, while A1 exhibited pronounced SSA for temporal configuration (Fig 5). This pattern challenges a purely feedforward adaptation account and instead supports a predictive-coding framework, in which higher-order cortical circuits detect deviations in temporal configuration and generate enhanced responses to deviants.

The weak SSA observed in response to click-train objects may reflect a general limitation in the SSA capacity of the IC neurons. This interpretation is supported by previous research showing that the IC typically exhibits only minimal stimulus-specific adaptation, even to pure tones [28,33,64]. By selecting neurons that showed strong SSA to complex harmonic tones, we confirmed that IC neurons are capable of robust SSA, yet this population still showed weak SSA for click-train configurations (S2 Fig). Thus, the lack of SSA to click-train configurations is not due to a general incapacity for SSA in IC neurons but reflects a specific difficulty in discriminating temporal configuration at this subcortical level.

Work with “tone clouds” provides an important comparison. Harpaz et al. (2021) used spectrally balanced tone clouds—each composed of the same set of tones presented once—and found early and strong SSA in A1 but only delayed and weak SSA in IC and MGB [62], suggesting that A1 encodes abstract temporal–spectral patterns. Tone clouds can be conceptualized as comprising a spectral configuration in the frequency domain and a temporal configuration in the time domain. Recent studies have shown that auditory cortical neurons can exhibit bursting responses selective for particular tone configurations rather than individual tones [65,66]. In contrast to these tone-based paradigms, we used identical pulses and removed spectral configuration, thereby demonstrating that purely temporal configuration is sufficient to drive discriminative responses in A1 neurons (Fig 5 and S2 Fig). Given the minimal SSA in IC and the robust SSA in A1 under these conditions, sensitivity to temporal configuration likely originates in higher auditory structures—MGB and especially A1—rather than in peripheral or early brainstem mechanisms.

Cooling experiments further support this view. Reversible suppression of auditory cortex substantially reduced configuration sensitivity in MGB (Fig 6), indicating that thalamic SSA for temporal configuration depends in large part on corticothalamic feedback. This is consistent with extensive reciprocal connectivity between MGB and auditory cortex [67] and supports a model in which cortical predictions and error signals shape thalamic responses to temporal structure.

Finally, layer-specific analysis revealed that supragranular layers of A1 exhibit stronger SSA for temporal configuration than infragranular layers, whereas SSA for complex tones did not show such pronounced laminar differences (Fig 7). This suggests that superficial layers—known to support horizontal integration and cortico–cortical communication—play a privileged role in encoding temporal configuration and in propagating prediction-error signals within the cortical hierarchy.

In summary, our findings highlight A1 as the principal site for integrating temporal configuration, with limited contributions from subcortical structures such as IC and MGB. This supports a hierarchical and predominantly top-down model of temporal processing in which cortical circuits detect deviations in fine temporal structure and feed predictions back to thalamus. Future work should identify the specific synaptic and circuit mechanisms—such as recurrent connectivity, inhibitory microcircuits, and corticothalamic loops—that implement sensitivity to complex temporal configurations and link these mechanisms to perception and behavior. In particular, relating neuronal sensitivity to temporal configuration to perceptual thresholds and biases in humans will further establish temporal configuration as a feature-like dimension of auditory representation.

## Methods

### Participants and Surgical Procedures

#### EEG Experiment Participants

We recruited 32 volunteers (18 males and 14 females; mean age = 25.3 years, SD = 3.59) with normal hearing to participate in the EEG experiments. Of these, 22 participated in Active Listening Session 1, 24 in Active Listening Session 2, and all 32 took part in the Passive Listening Sessions. The study protocol adhered to ethical standards and was approved by the Institutional Review Board (IRB-20230131-R). Informed consent was obtained from all participants prior to participation.

#### EEG Experimental Procedures

The study comprised six experimental sessions, including two active discrimination sessions and four passive listening sessions.

##### Active Listening Session 1 (Fig 1B)

In this session, participants engaged in a delayed match-to-sample task. An initial click train (one of the ten click trains listed in S1 Table) was played, followed by a 500-ms silence before presenting a second click train, randomly selected from the same pool of ten options. Participants were required to assess whether the two sounds were identical. Responses were recorded using the left arrow key for differing sounds and the right arrow key for identical sounds. We selected 20 different sound pairs, as shown in Fig 1C, each presented 40 times per session. Of these, 7 pairs involved identical sounds, while 13 pairs consisted of different sounds.

##### Passive Listening Sessions 2-5

Following the two active discrimination sessions, four passive listening sessions were conducted. During these sessions, participants were exposed to auditory stimuli without performing any behavioral tasks. Each session consisted of a continuous sequence of auditory stimuli, with each sound lasting 200 ms and presented consecutively with a 300-ms interval between the onset of consecutive sounds, totaling 2400 presentations. The stimuli included seven click trains randomly selected from a pool of ten different click trains. One click train was presented most frequently (2160 times) and served as the standard stimulus, while the other six types were used as deviant stimuli, each presented 40 times and interspersed among the standard sounds. The configuration ensured that each deviant stimulus was preceded by 7 to 11 standard stimuli, preventing consecutive presentations of deviant stimuli.

- Session 3: Standard stimulus was ‘Con’; deviant stimuli included ‘Asc’, ‘RN_10’_, ‘RN_40’_, ‘RN_60’_, ‘TW_30’_, and ‘TW_50’_.
- Session 4 (Fig 2A): Standard stimulus was ‘TW_30’_; deviant stimuli included ‘Asc’, ‘RN_10’_, ‘RN_40’_, ‘RN_60’_, ‘TW_10’_, and ‘TW_50’_.
- Session 5: Standard stimulus was ‘RN_60’_; deviant stimuli included ‘Con’, ‘RN_10’_, ‘TW_10’_, ‘TW_30’_, ‘TW_50’_, and ‘TW_70’_.
- Session 6: Standard stimulus was ‘RN_10’_; deviant stimuli included ‘Con’, ‘Asc’, ‘RN_40’_, ‘TW_10’_, ‘TW_30’_, and ‘TW_50’_.

Participants were instructed to maintain a stationary head position throughout the experiment and were allowed a 5-minute break after each session to prevent fatigue and ensure attentiveness.

#### ECoG surgery

Three adult male Wistar rats (280 - 340 g, 9 - 12 weeks), each with clean external ears, were selected for this study. We implemented sterile surgical techniques for the implantation of the headpost and the ECoG array. Anesthesia was administered using pentobarbital sodium (40 mg/kg) along with atropine sulfate (0.05 mg/kg, s.c.) given 15 minutes before the procedure to reduce tracheal secretion. Xylocaine (2%) was applied liberally to the incision site to minimize pain. A head fixation bar was secured to the top of the skull using dental cement and six titanium screws. A craniotomy exposed a 4.5 mm × 5 mm area above the auditory cortex on the left side, leaving the dura mater intact. The ECoG array (KD-PIFE, KedouBC, China), consisting of 32 electrodes in a 6×6 grid with 1 mm spacing, was implanted. The array covered an approximate area of 5 mm ×5 mm, encompassing the entire auditory cortex (AC) (Fig 3A). After the implantation, the ECoG array were covered with artificial brain gel, followed by the repositioning of the bone flap over the surgical site. The area was then sealed with dental cement to ensure closure and protection. Postoperative recovery lasted 7 days, during which anti-edema medications and antibiotics were administered, and the animals’ weight was monitored daily to ensure their well-being. This component of the study received approval from the Animal Subjects Ethics Committees of Zhejiang University (ZJU20210078).

#### Extracellular recording surgery

Three adult male Wistar rats (280 - 340 g, 9 - 12 weeks) were utilized for extracellular electrode recordings, undergoing a similar procedure to the ECoG surgery for head post stabilization. Then, the left lateral and dorsal surfaces of the skull were exposed post-surgery. Once cleaned and dried, reference landmarks were established at the bregma and on the left lateral skull for precise electrode positioning. Target areas for the electrode recordings included the left primary auditory cortex, the left medial geniculate body, and the left inferior colliculus (IC). To prevent tissue overgrowth, the skull above these areas was thinly coated with dental cement and enclosed with a protective wall. A handheld micro-drill created a small opening (approximately 2 mm × 2 mm) at the designated target site, which was subsequently sealed with brain gel. On the recording day, the dura mater was carefully punctured, and the electrode was vertically inserted into the target area using the established landmarks for guidance.

#### Cooling AC

To investigate the functional role of AC in processing click-train temporal configurations, we employed a reversible cooling technique to deactivate the AC and recorded activity from the ipsilateral medial geniculate body (MGB). A 4-mm diameter cryoloop was positioned over the AC, manually bent to fit the cortical surface and covering both the primary and non-primary fields. Cooling was achieved by circulating anhydrous ethanol, cooled by dry ice, through the cryoloop using a peristaltic pump. A temperature probe was placed near the AC to continuously monitor the temperature, which was regulated by adjusting the ethanol flow rate, maintaining a stable temperature of 8 ± 3°C. This cooling temperature effectively deactivates all layers of the AC, including corticofugal projections from layers V and VI, without significantly affecting neural activity in subcortical areas such as the MGB [67]. MGB neuronal activity was recorded 5 minutes after cooling began. Additionally, we monitored AC neuron activity before, during, and after cooling to confirm the deactivation effect and ensure that the structural and functional integrity of the deactivated area remained intact.

### Auditory stimuli and Experimental Procedures

#### Stimuli

We developed three experimental procedures for EEG, ECoG, and extracellular recording, employing ten unique click trains as depicted in S1 Table. Each click train lasted 200 ms, consisting of 51 fixed pulses, with each pulse lasting 200 µs and an average inter-click interval (ICI) of 4 ms. The sound intensity for all click trains was uniformly calibrated to 60 dB SPL to ensure consistency across experiments involving both human participants and animal subjects. Calibration was performed using a ¼-inch condenser microphone (Brüel & Kjær 4954, Nærum, Denmark) and a PHOTON/RT analyzer (Brüel & Kjær, Nærum, Denmark).

For human EEG recordings, click sounds were synthesized using MATLAB R2022b (The MathWorks, Inc., Natick, MA, USA) on a computer equipped with a high-fidelity sound card (Creative, AE-7). These clicks were produced with a sampling rate of 384 kHz and 32-bit precision and then broadcast through a stereo speaker system (Golden Field M23), ensuring the precise presentation of click train sequences. In contrast, animal ECoG and Extracellular recording experiments were conducted within a sound-proof room, with stimuli played contralaterally to the recording site in rats. These acoustic stimuli were digitally generated using a computer-controlled Auditory Workstation (RZ6, TDT) at a sampling rate of 97656 Hz and delivered through magnetic speakers (MF1, TDT) within the TDT systems.

#### ECoG experimental procedures

Four passive listening sessions, identical to those used in human EEG studies, were adapted for use with rats. These sessions involved playback of the same auditory stimuli configured for the human experiments.

#### Extracellular recording experimental procedures

The study included six sessions specifically designed for extracellular recordings.

*Session 1 Frequency screening procedure*. A sequence of tones was presented randomly at the contralateral position to the recording site. Each tone lasted 100 ms with a rise-fall time of 5 ms. The frequency range spanned from 0.5 kHz to 48 kHz, divided into 26 logarithmic steps. The intensity of these tones varied from 0 to 70 dB SPL in increments of 10 dB. Each combination of frequency and intensity was repeated five times with a 300-ms interval between stimuli. This session aimed to measure basic neuronal response properties, specifically the frequency response area (FRA) and characteristic frequency (CF).

*Session 2-4 Oddball procedure*. We selected two click trains from the set of ten click trains mentioned earlier to create a stimulus pair, thereby establishing classic oddball sequences (refer to Fig 4B). Each sequence of click trains comprised 300 presentations with an inter-stimulus interval (ISI) of 250 ms. The selected pair of click trains was presented in two separate oddball sequences and 9 standard stimuli were followed by 1 deviant stimulus in each sequence. Hence, the standard sequence, which included 270 out of 300 presentations (90%), and the deviant sequence, which included 30 out of 300 presentations (10%). Each of the two click trains was used once as a standard and once as a deviant stimulus. The key distinction among *Session 2*, *Session 3* and *Session 4* lies in the different stimulus pairs utilized, while all other experimental parameters remained constant. Specifically, for *Session 2*, the stimulus pair comprised ‘RN_10’_ and ‘TW_50’_; for *Session 3*, the stimulus pair comprised ‘Asc’ and ‘TW_30’_, whereas for *Session 4*, it consisted of ‘RN_60’_ and ‘Con’.

*Session 5 Complex Tone Oddball procedure*. To complement the findings obtained from click train pairs, we designed an oddball sequence utilizing two complex harmonic tones. One of the complex harmonic tones comprised frequencies with intervals of 1/8 octaves per step, starting at a fundamental frequency of 600Hz. The harmonic frequencies (654 Hz, 712 Hz, 777 Hz, …, 39910 Hz) followed this progression. The other complex harmonic tone started at a fundamental frequency of 680 Hz, also using 1/8 octaves per step. The two complex harmonic tones did not share similar frequencies. Except for the frequency components, other stimulus parameters for this oddball procedure were identical to those employed for the click trains.

### Data collection and pre-processing

#### EEG recording

EEG recording was conducted using a 64-channel electrode cap (10-20 system, NeuSen W series, Neuracle, China) connected to a wireless amplifier (NeuSen W series, Neuracle, China). Of the 64 channels in the electrode cap, five electrodes (’ECG’, ‘HEOR’, ‘HEOL’, ‘VEOU’, ‘VEOL’) were dedicated to recording electrooculography (EOG) and electrocardiogram (ECG), leaving 59 channels (refer to Fig 2B) for processing signals. Additionally, two electrodes positioned slightly behind and in front of the Fz electrode served as the reference electrode (REF) and the ground electrode (GND), respectively, for referencing and grounding purposes. EEG data and the time of trial onset were simultaneously recorded at a sampling rate of 1 kHz. Participants were comfortably positioned with head support in an acoustically isolated room during the experimental sessions. They were instructed to minimize eye blinks and facial movements to reduce artifacts.

The raw EEG data were band-pass filtered between 0.5 Hz and 40 Hz and epoched based on the deviant stimuli or the standard stimuli preceding the deviant trigger times, encompassing 750 ms before and 1000 ms after the trigger, respectively. Baseline correction was then applied by subtracting the mean responses within the rest signals before the sequence start. To identify and reject motion artifacts, a relative threshold approach was utilized for each epoch. Epochs containing data points exceeding predefined thresholds were excluded. The relative threshold was determined based on the percentage of bad samples within a trial. A sample was flagged as a “bad sample” if it fell outside the range of “Mean” ± 3 × “SD” for a trial (epoch) across specific channels. Trials with over 20% of bad samples were categorized as “bad trials”. Additionally, channels with over 10% of bad trials were marked as “bad channels”. Initially, bad channels were excluded by evaluating bad samples using data from all channels. Subsequently, bad trials were excluded from all channels by computing bad samples using data from the remaining good channels. Finally, independent component analysis (ICA) was employed using FieldTrip Toolbox [68] to remove additional artifacts, specifically those related to EOG and ECG.

#### ECoG recording

ECoG recording involved positioning a reference wire from the ECoG array into the subdural space, with the ground wire connected to a titanium screw at the front of the skull. Lead wires from the ECoG array were then connected to micro connectors (ZIF-Clip headstage adapters, Tucker-Davis Technologies, TDT, Alachua, FL). The signals were subsequently amplified using an amplifier (RZ5, TDT), digitally sampled at a rate of 12 kHz, and stored on hard drives for later detailed analysis.

The raw ECoG signal underwent down-sampling to 600 Hz and was segmented into epochs lasting 1.75 seconds each (comprising 0.75 seconds before the deviant or standard stimulus and 1 second after). Subsequently, baseline correction was applied, and any identified bad channels or trials were excluded from further analysis. The pre-processing methodology mirrored that used for EEG data.

#### Extracellular recording

Extracellular recording involved the use of Version 3A Neuropixels silicon probes (Imec, Leuven, Belgium; https://github.com/cortex-lab/neuropixels/wiki/About_Neuropixels) during each session. Each puncture was precisely positioned based on pre-established markers. A flexible ground wire was soldered to the electrode’s PCB, with the ground and reference contacts shorted. Throughout the experiments, this ground wire was connected to a titanium screw connector at the front of the skull. The probes were cautiously inserted vertically through 0.3−0.5 mm long slots in the dura, avoiding visible blood vessels to minimize damage to brain tissue. A single-axis motorized micromanipulator, controlled from outside the soundproofed room, facilitated the insertion process. The average insertion depth for A1 was approximately 2 ±0.1 mm (mean ±SD, n = 12), MGB: 7 ±0.5 mm (n = 14), IC: 6.1 ± 0.4 mm (n = 12) below the brain surface, in accordance with a standard rat brain atlas[69]. White noise stimuli were employed to ascertain the presence of auditory responses.

The signals were recorded using the Neuropixels PXIe control system (Imec, Leuven, Belgium) at a sampling rate of 30 kHz for spike data and 2.5 kHz for local field potentials (LFPs). Neuronal spike trains were sorted offline using the Kilosort algorithm, followed by manual curation in Phy [70]. Subsequently, for each neuron, the response window (0–100 ms) and baseline window (–20–0 ms) were defined relative to the start of each oddball sequence. As the response and baseline windows were not equal in duration, spike counts in each window were converted to firing rates (spikes per second) prior to analysis. We also calculated the poisson cumulative distribution function from the baseline to assess the likelihood that the observed spike counts at each time point could occur randomly [71], and determined the latency as the time point of the first spike where the Poisson probability falls below the specified threshold (1e-3). Neurons were classified as responsive according to three criteria: 1) the firing rate in the response window significantly exceeded that in the baseline window (right-tailed paired t-test, α = 0.05); 2) the response latency could be detected; 3) the average spike count in the response window had to be at least one spike higher than that in the baseline window. Neurons meeting these criteria were defined as auditory-responsive and were utilized for subsequent analytical calculations. For LFPs, we downsampled the signals to 300 Hz for subsequent current source density (CSD) analysis.

All recordings of AC data utilized for subsequent analysis were conducted exclusively from the A1 in the left hemisphere, identified by the tonotopic gradient of CF. Additionally, upon completion of the entire recording session, the Neuropixel probes were coated with DiI and inserted into the MGB and IC areas for precise positioning [62]. Subsequently, the rat underwent transcardial perfusion in a fume hood. An example of the recording location in the rat cortex is illustrated in Fig 4D-4E.

In total, recordings were obtained from 671 neurons in the A1, 632 neurons in the MGB, and 940 neurons in the IC. All recorded neurons were included in the subsequent analysis. Furthermore, a subset of these neurons underwent control experiments utilizing complex harmonic tone oddball paradigms (*Session 5* in Extracellular recording experimental procedures). From this subset, to assess the statistical significance of the CSI for each neuron, we employed a nonparametric bootstrap procedure with 1,000 iterations to generate a null distribution. We then evaluated whether zero fell outside the lower bound of the 95% confidence interval (CI), indicating that the CSI was significantly greater than zero. Only neurons with CSI values significantly greater than zero were included in subsequent analyses. This procedure included 94 neurons in A1, 109 neurons in MGB, and 140 neurons in IC.

### Data Analysis

All steps of the mentioned off-line EEG, ECoG and Extrcellular data processing were performed using MATLAB R2022b and the FieldTrip toolbox [68].

#### EEG data analysis

EEG data analysis involved computing the relative response magnitude to quantify the difference between the responses to deviant and standard stimuli, utilizing average evoked response potential (ERP) data from each channel across trials for each subject. For individual subjects (refer to Fig 2D-2E), we selected the 0-300 ms window relative to the onset of both deviant and standard stimuli as the calculation window. A grand-average ERP waveform was then obtained by averaging data across selected interesting channels from each subject. These channels were situated on the temporal, parietal, and occipital lobes. In total, 27 channels were utilized in following processing.

Furthermore, we computed the mean amplitude of the average response from each channel. For this computation, we applied a sliding window approach with a window size and step size of 5 ms. Subsequently, an independent-samples t-test with FDR correction was conducted to determine if there was a significant difference between the mean values of the deviant and standard conditions across channels.

For population analysis, we averaged the grand-average waveform across all subjects (refer to Fig 2F-2I). We also calculated the mean value on every grand-averaged waveform using the same sliding window parameters and conducted an independent-samples t-test with FDR correction to assess the differences between the deviant and standard stimulus conditions across subjects. If the p-value for the difference between the deviant and standard stimuli is less than 0.05 during this time period, the corresponding time points are marked in grey.

#### ECoG data analysis

For each rat, we computed the average ERP from each channel across all trials. Using the same time window employed for EEG data analysis, we calculated the amplitudes for both deviant and standard stimuli. Subsequently, paired t-tests were conducted to evaluate whether there were significant differences between the responses to deviant and standard stimuli on each channel. Specifically, time points where the p-value for the difference between deviant and standard stimuli was less than 0.05 during this period were marked in grey.

For population-level analysis, we calculated the mean amplitude of the ECoG waveforms within a 0–200 ms window following the onset of each deviant stimulus and compared it with the mean amplitude during a 50 ms baseline window preceding the stimulus. Channels from each rat where the response exceeded 2 standard deviations above the baseline were selected and pooled together (N = 3 rats, n = 36 channels). These pooled channels were averaged to create a population-level waveform. In this combined dataset, we followed the same statistical testing approach described above to determine time periods where significant differences existed between deviant and standard stimuli. Time points where the p-value for the difference between deviant and standard stimuli was less than 0.05 during this period were marked in grey.

#### Extralcellular data analysis

For oddball paradigms, we employed one SSA indice consistent with previously published work [31,42,72,73] to describe the adaptive degree of SSA.

The Common-SSA Index (CSI), was computed using the formula:

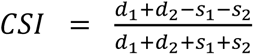

Here, s_1 a_nd d_1 r_epresent the neuronal responses to the first click train in an oddball pair when it was the standard and deviant, respectively, while s_2 a_nd d_2 d_enote the responses to the second click trian in the pair under the same conditions. The SSA index CSI equals 1 when adaptation is complete (i.e., no response to the standard, and significant response to the deviant), and 0 when there is no adaptation (i.e., the response to the standard and deviant is equal). Meanwhile, to assess the significance of the CSI for each unit, we employed a nonparametric bootstrap procedure with 1,000 iterations to generate a null distribution of the CSI values. The 95% confidence interval (CI) of the bootstrapped distribution was then computed. A unit was considered to have a significantly greater-than-zero CSI if the lower bound of the 95% CI exceeded zero. Neurons passing this criterion were highlighted in red to indicate their distribution in the corresponding figure.

Following spike count computation, the window was consistently chosen from 5 to 200 ms relative to the onset of the deviant stimuli or the ninth standard stimuli. It should be noted that all data analyzed using ANOVA or t-test in this study were assessed for normality using the Shapiro-Wilk test prior to analysis. Otherwise, the Wilcoxon signed-rank test or Mann-Whitney U test was used.

For all peri-stimulus time histograms (PSTHs), the bin size was set to 5 ms, step was set to 5ms. To generate the PSTH heatmap (Fig 5), the neural responses of the population to each deviant and standard stimulus were quantified and normalized using z-score transformation. The z-scored responses to standard stimuli were subtracted from those to deviant stimuli for each neuron. The resulting difference values were then used to construct the heatmap, where each row represents an individual neuron and the color scale reflects the z-scored difference.

#### Current Source Density Analysis

One-dimensional current source density (CSD) profiles were calculated from the second spatial derivative of the LFP [40,74–76], which can be approximated using the following formula:

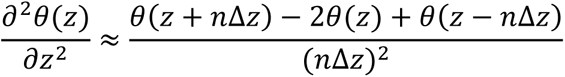

where θ is the field potential, z is the spatial coordinate perpendicular to the cortical laminae, Δz is the spatial sampling interval (Δz = 80 µm), and n is the differentiation grid (n = 2). To reduce spatial noise, a five-point Hamming filter was applied [75]. The LFP data used for this analysis were selected from channels with an 80 µm sampling interval, as we used a dense channel configuration with Neuropixel electrodes (example shown in S4 Fig, left). The sound stimulus was a 100-ms noise burst. In the resulting laminar CSD profiles, current sinks are shown in red and current sources in blue (S4 Fig, right). Visualization of laminar profiles was enhanced using linear channel interpolation.

In this text, we describe and discuss our A1 results in terms of cortical layers. Our divided approach is primarily based on two criteria: first, we compared the electrode penetration depth with previously published depth information of cortical layers [77,78]; second, we utilized CSD profiles. Specifically, our CSD sink latency depth profiles showed excellent agreement with those previous studies [40,75,76,79], from which we adopted their layer classification methodology. Based on these two criteria, we divided the cortical layers into three parts (supra: layer I/II, middle: layer III/IV and deep: layer V/VI) for our analysis.

## Acknowledgments

We are grateful to Profs. Xiaowei Chen and Xi Chen for their invaluable comments on the early version of the manuscript, as well as to Xiaokai Kou for the help with the experiments.

## Supporting information

**S1 Fig.**
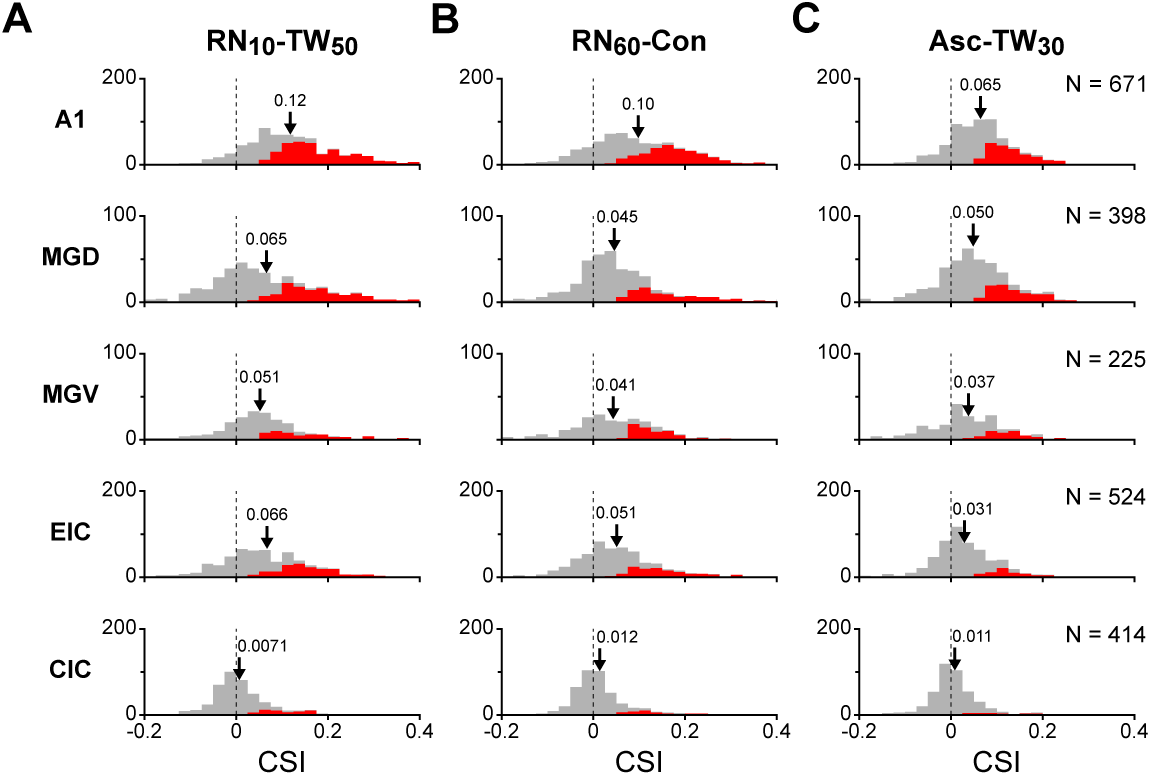
Populational neuronal response to oddball paradigm with temporally merged objects in A1 and subregions of IC, MGB. **(A)** Distribution of CSI in A1 (n = 671), MGV (n = 398), MGD (n = 225), CIC (n = 524) and EIC (n = 414) in the oddball paradigm (RN_10,_ TW_50)_. Arrows indicate the mean. Red bars highlight neurons with CSI significant greater than 0, determined by a nonparametric bootstrap procedure with 1,000 iterations (p < 0.05). **(B-C)** Distribution of CSI values to additional two sound pairs in the oddball paradigm (RN60, Con) and (Asc, TW_30)_.

**S2 Fig.**
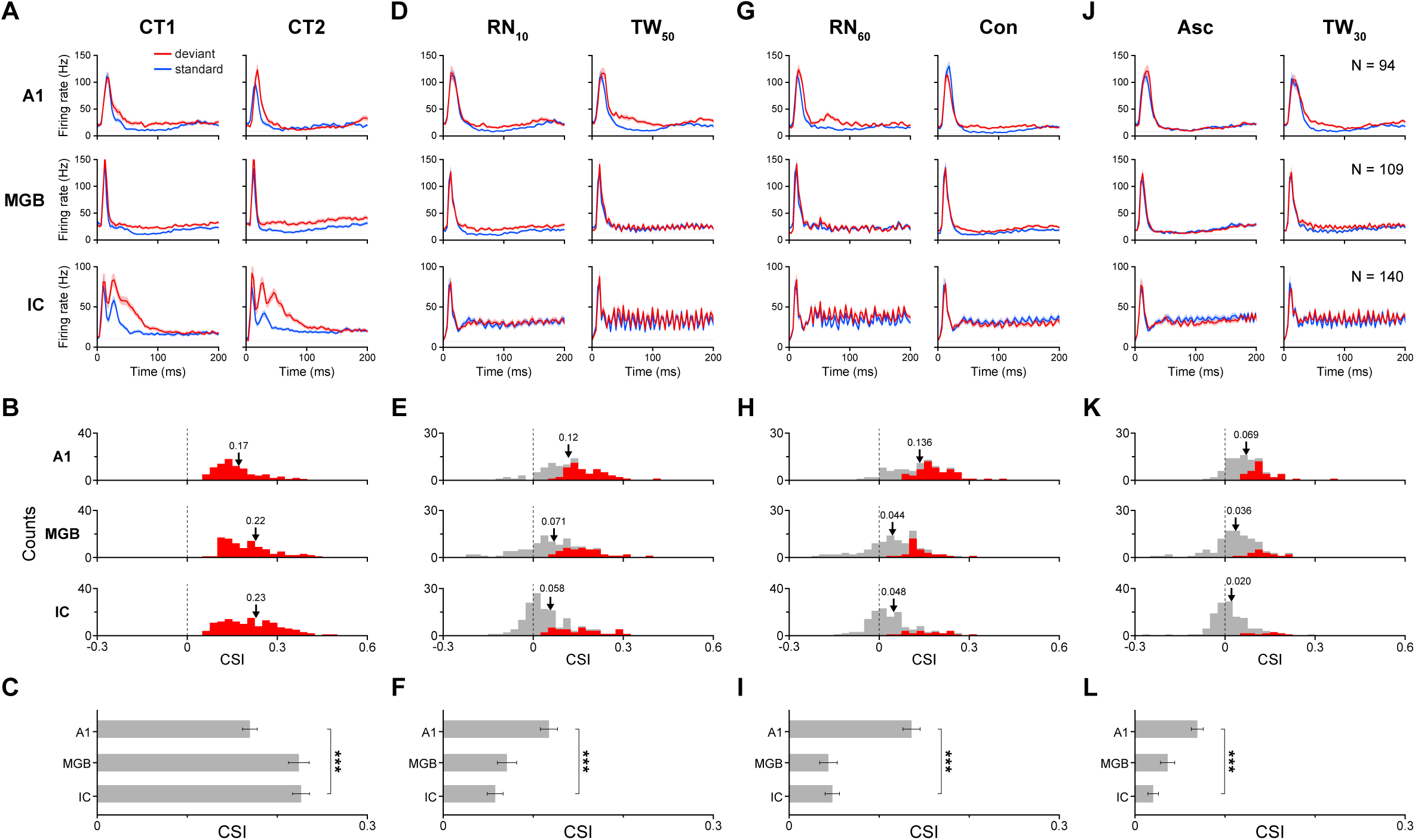
Populational neuronal responses to complex tones and temporally merged objects in the oddball paradigm. **(A)** Populational PSTHs displaying responses to two complex tones as deviant (red) and standard (blue) stimuli across three auditory stations: A1 (n = 95, top row), MGB (n = 109, middle row), and IC (n = 140, bottom row). Only neurons with significant CSI values, determined by a nonparametric bootstrap procedure with 1,000 iterations (p < 0.05), were included in these datasets. **(B)** Distribution of CSI in A1 (top row), MGB (middle row), and IC (bottom row). Arrows point to the mean. Red bars highlight neurons with CSI significant greater than 0, determined by a nonparametric bootstrap procedure with 1,000 iterations (p < 0.05). **(C)** Comparison of CSI among A1, MGB, and IC. Error bars represent standard error. ***, p < 0.001, ANOVA with post-hoc test. **(D-F)** Populational responses from the same datasets featured in panels (A-C) to the sound pair in the oddball paradigm (RN_10,_ TW_50)_: Adheres to the same conventions as panels (A-C). **(G-I)** Populational responses from the same datasets featured in panels (A-C) to the sound pair in the oddball paradigm (RN_60,_ Con): Adheres to the same conventions as panels (A-C). **(J-L)** Populational responses from the same datasets featured in panels (A-C) to the sound pair in the oddball paradigm (Asc, TW_30)_: Adheres to the same conventions as panels (A-C).

**S3 Fig.**
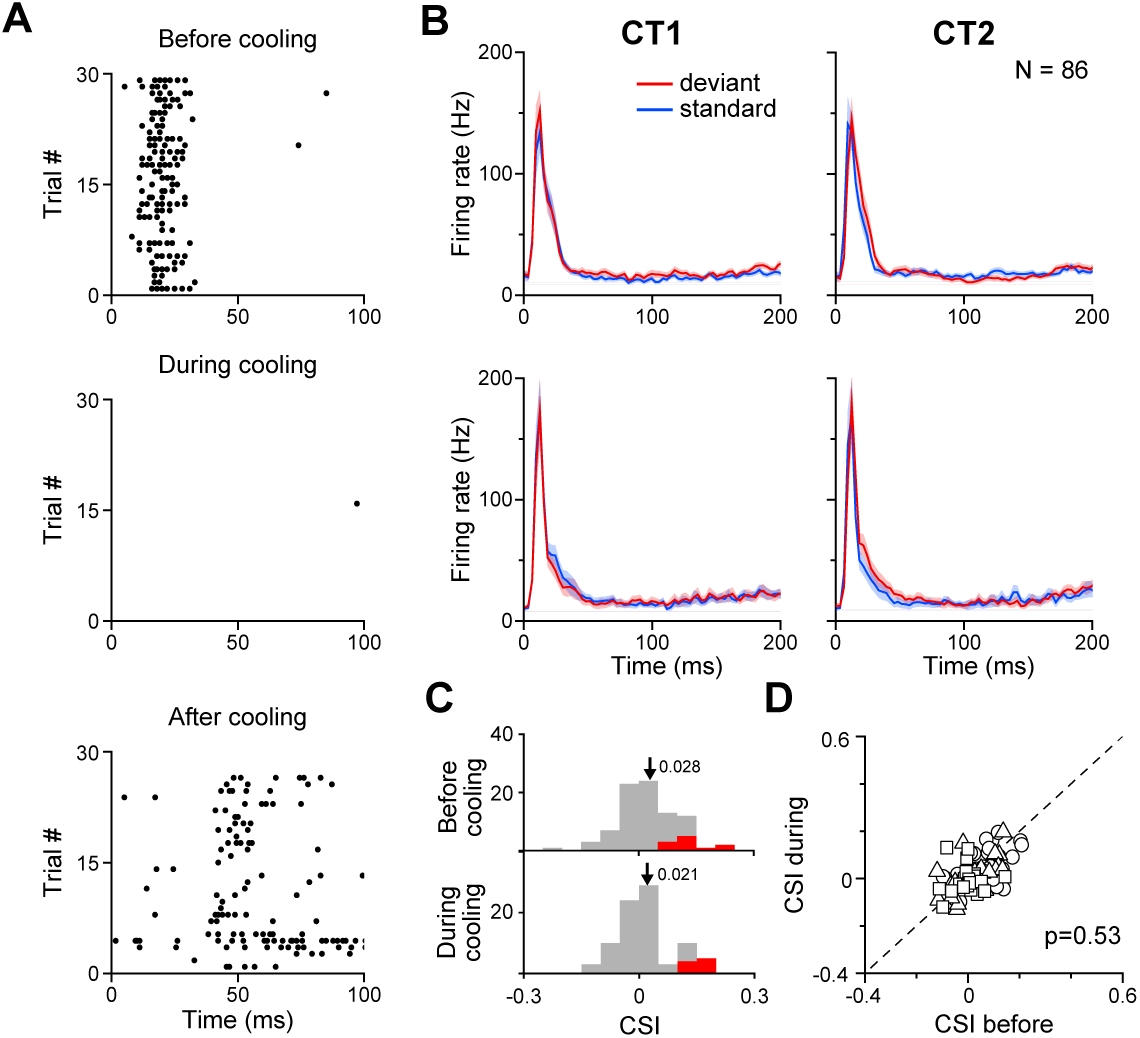
Effect of AC cooling on the response in A1 and oddball paradigm with complex tones in MGB. **(A)** Neuronal responses in the AC before, during, and after AC cooling. Raster plots display the responses to the noise sound (duration = 100 ms) in each of the three conditions: before cooling (top row), during cooling (middle row), 2 hours after cooling (bottom row). **(B-D)** Populational responses to two complex tones in the oddball paradigm, adhering to the same conventions as Fig. 6, p=0.53, paired t-test.

**S4 Fig.**
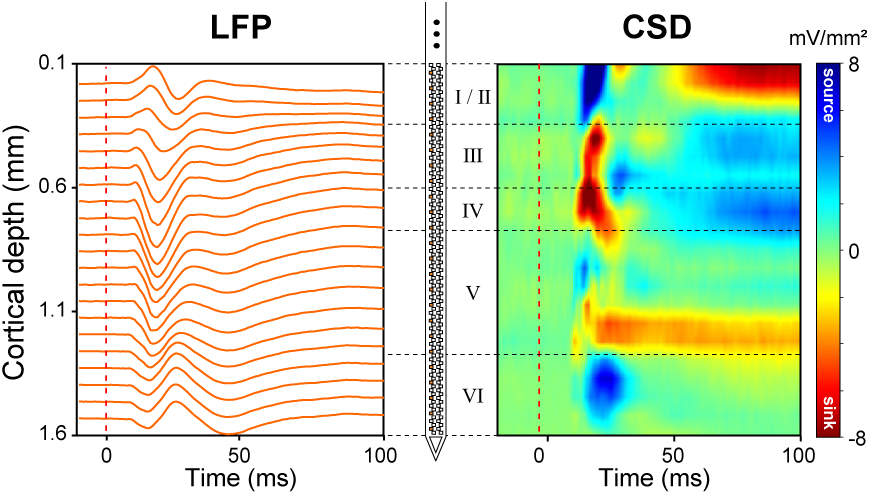
Laminar profiles of evoked responses to noise recorded from a single electrode penetration into A1. A schematic of the 384-channel neuropixel electrode is shown in the center of the figure. The electrode records local field potentials (LFPs, left) from superficial to deep layers, with specific channels selected every 80 µm (marked in orange) for analysis. The right panel shows current source density (CSD), calculated from the LFP values of the selected channels and smoothed for better viewing. For visualization, the response amplitude is color-coded, with sources in blue and sinks in red. Horizontal dashed lines represent the rough boundaries of cortical layers. Noise onset times are indicated by vertical dashed lines.

**S1 Table.** Description of the ten sets of click trains.

