## Supplementary material for "Hierarchical Encoding of Millisecond-Scale Temporal Configuration of Sound in the Mammalian Auditory Pathway": S1 Table

| Num | Name | Average ICI (ms) | Pulse Num | Description |
|---|---|---|---|---|
| 1 | Con | 4 | 51 | ICI remains constant at 4ms |
| 2 | Asc | 4 | 51 | ICI increases progressively from 2 ms to 6 ms |
| 3 | Dec | 4 | 51 | ICI decreases progressively from 6 ms to 2 ms |
| 4 | $TW_{10}$ | 4 | 51 | ICI varies randomly between two values: 3.6 ms and 4.4 ms (4±10%) |
| 5 | $TW_{30}$ | 4 | 51 | ICI varies randomly between two values: 2.8 ms and 5.2 ms (4±30%) |
| 6 | $TW_{50}$ | 4 | 51 | ICI varies randomly between two values: 2 ms and 6 ms (4±50%) |
| 7 | $TW_{70}$ | 4 | 51 | ICI varies randomly between two values: 1.2 ms and 6.8 ms (4±70%) |
| 8 | $RN_{10}$ | 4 | 51 | ICI varies randomly within the range of 3.6 ms to 4.4 ms (4±10%) |
| 9 | $RN_{40}$ | 4 | 51 | ICI varies randomly within the range of 2.4 ms to 5.6 ms (4±40%) |
| 10 | $RN_{60}$ | 4 | 51 | ICI varies randomly within the range of 1.6 ms to 6.4 ms (4±60%) |
